# Organ-specific translation elongation rates measured by in vivo ribosome profiling

**DOI:** 10.1101/279257

**Authors:** Maxim V. Gerashchenko, Zalan Peterfi, Vadim N. Gladyshev

**Affiliations:** Division of Genetics, Department of Medicine, Brigham and Women’s Hospital, Harvard Medical School, Boston, MA, USA

**Keywords:** protein synthesis, translation, elongation, ribosome profiling, mouse

## Abstract

Protein synthesis and degradation are intricate biological processes involving more than a hundred proteins operating in a highly orchestrated fashion. Despite the progress, few options are available to access translation in live animals as the increase in animal’s complexity limits the repertoire of experimental tools that could be applied to observe and manipulate processes within animal’s body, organs, and individual cells. It this study, we developed a labeling-free method for measuring organ- and cell-type specific translation elongation rates. It is based on a time-resolved delivery of translation initiation and elongation inhibitors in live animals followed by ribosome profiling. It also reports translation initiation sites in an organ-specific manner. Using this method, we found that the elongation rates differ among mouse organs and determined them to be 6.8, 5.2, and 4.4 amino acids per sec for liver, kidney, and skeletal muscle, respectively.

**Significance:** Protein synthesis is a vital biological process. Modern methods of genome editing enable generation of sophisticated animal models to study the regulation of protein synthesis in health end disease. However, the methods that could track various steps of translation at a gene level resolution *in vivo* are lacking, particularly in complex vertebrates, such as mice and rats. Here, we measured the translation elongation rate in several organs by delivering inhibitors specific to certain phases of translation directly through the mouse bloodstream. This study lays out a path for interrogating translation in animals in response to various genetic and dietary interventions.

---

Protein synthesis is regulated both at the transcriptional (availability of mRNA) and translational (the number of active ribosomes, translational efficiency, the speed of elongation and the frequency of initiation events) levels. This complexity presents endless possibilities for malfunctioning. In turn, the vast majority of cellular functions are carried by proteins, making any aberrations in their biosynthesis potentially harmful. Modern *in vitro* methods allow unsurpassed precision and flexibility, e.g. real-time translation dynamics of a single molecule can be monitored with fluorescent microscopy (1, 2), and transcriptome-wide snapshots of translation with a single nucleotide resolution can be achieved by means of ribosome profiling (3–6). In contrast to research performed in cells, few options are currently available to investigate translation in live animals. It is particularly challenging to assay specific steps of translation, i.e. elongation, initiation or termination rate, *in vivo*. Several clever approaches were designed, including animals fed with amino acid supplements labeled with a stable isotope (7) or radioactively labeled amino acids. To estimate the average translation elongation speed, short-term pulses of radioactively labeled amino acids can be injected directly into the portal vein of an animal (8). The radiation accumulation rate can be used to project the average translation elongation rate (8, 9); however, tissue samples are rendered by this method unsuitable for many other molecular biology assays.

We were guided by these challenges and the need to assess various steps of translation in various organs of live animals at an individual gene level. In this work, we present a technique to directly assess translation *in vivo* and measure organ- and cell-specific translation elongation rates and other features of protein synthesis. Our method requires no radioactive labeling or transgenic reporters and can be applied to a variety of small animals such as a mouse or a rat. It relies on two translation inhibitors injected directly into the bloodstream in a time-dependent fashion. The first inhibitor (harringtonine) blocks initiation of translation without affecting elongation. The second inhibitor (cycloheximide) is injected after a specified time (less than a minute) to block elongation. Thus, the time-dependent run-off of ribosomes is detected after the analysis of sequencing data and the elongation rate and other features of translation are inferred.

## Results

### Translation inhibitors perform well in live mice

We first determined whether two translation initiation inhibitors known to work in cell culture - lactimidomycin (10) and harringtonine (11) – could affect translation upon injection in the blood-stream of a mouse, and used C57BL/6 males (13-15 week-old) as an experimental cohort. Drug delivery was performed via the retroorbital injection (12). These inhibitors are expected to cause depletion of the polysomal fraction, which can be assessed by sucrose gradient fractionation of organ lysates (Fig. 1A-B). Within 5 min, harringtonine (0.5-4 mg/mouse) completely depleted polysomes in all organs except for testes and brain, where up to 30 min were needed to significantly reduce the number of polysomes due to the blood-brain and blood-testes barriers. Harringtonine specificity was further confirmed by sequencing mRNA fragments trapped within the ribosome active core, the method also known as ribosome profiling or Ribo-seq. Post-treatment ribosomes were almost exclusively located near the start codons, agreeing with the notion that harringtonine primarily affects initiation of translation (Fig. 1C). During these experiments, we noticed that the high doses of harringtonine may cause cardiac arrest in some animals; therefore, we adjusted the dosage and monitored the heart function with the electrocardiogram (ECG, Fig. 2B-C). The optimal dose was found to be 200 *μ*l of harringtonine (2.5 mg/ml) in the phosphate buffer saline per 20-25 g animal (Fig. 2B). Next, we co-injected harringtonine with varying doses of cycloheximide (20-60 mg/ml) to test the efficacy of the latter. When administered together, these drugs should not have a noticeable effect on the polysome profile, which is exactly what we observed (Fig. 1A, Fig. S1A). Hence, both inhibitors were active *in vivo* and had comparable uptake rates across tissues.

**Fig. 1.**
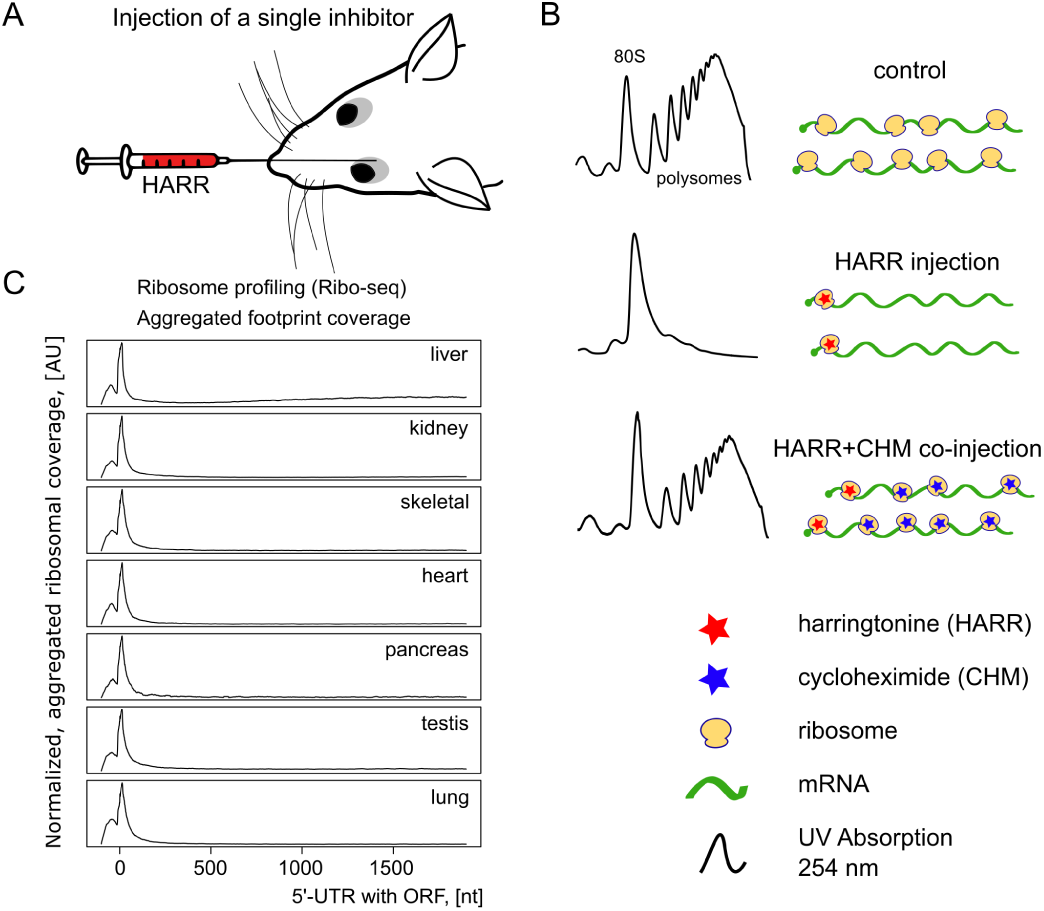
Evaluating translation inhibitors *in vivo*. (A) Single injection done to retroorbital cavity around the eye. (B) Both translation inhibitors show similar efficiencies and uptake times following injection into the bloodstream. Using sucrose gradient profiles of the mouse liver as an example, we demonstrate that harringtonine depletes polysomes while co-injection with cycloheximide preserves polysomes. (C) Harringtonine performance in 7 organs. It effectively stalls ribosomes at the vicinity of start codons. Post-injection incubation time was 5 min except testes, where up to 30 min were required to pass the blood-tissue barriers. Zero corresponds to the start of the open reading frame. 100 nucleotides from the 5’-UTR were added to allow mapping footprints over the start codon.

**Fig. 2.**
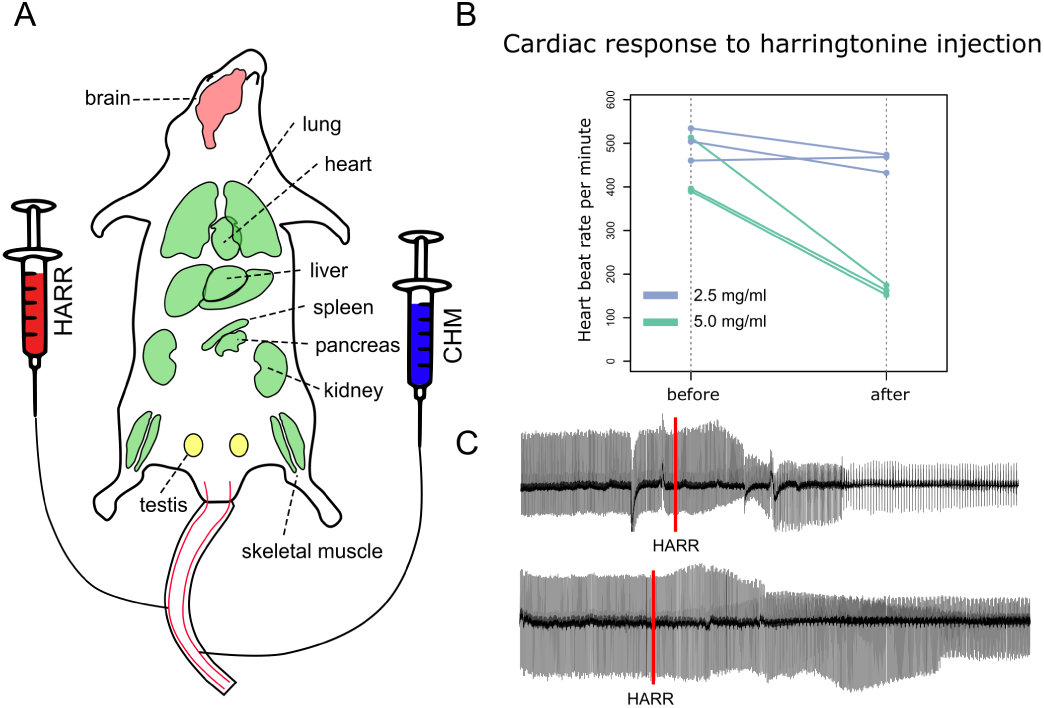
Experimental setup. (A) Layout of mouse organs susceptible to translation inhibitors. Organs marked in green are permeable for both harringtonine and cy-cloheximide within 15 sec post-injection. Translation elongation rate can be directly measured in these organs. Testes (shown in yellow) are partially permeable, and brain is mostly unaffected by inhibitors. (B) High doses of harringtonine severely drop heartbeat rate. We established that 2.5 mg/ml had no effect on the beat rate, whereas 5 mg/ml decreased it twice. 10 mg/ml caused cardiac arrest in most of the cases. All injections were done in a tail vein - injection volume was 200 *μ*l and the weight of mice was 20-25 g. (C) Representative electrocardiograms (ECG) from injected mice. The upper track shows the slowdown of the beat rate in response to 5 mg/ml harringtonine. The lower track shows no difference after the injection of 2.5 mg/ml harringtonine. The effect of harringtonine appears roughly 20 sec after the injection. The red line marks the injection event.

We also followed on some reports suggesting that translation can be affected by volatile anesthetics such is isoflurane used to sedate and put mice asleep (13). No evidence of isoflurane impacting translation was observed under our experimental conditions (Fig. S2).

## Two translation inhibitors can be used to produce a time-dependent shift in ribosome occupancy

Given the two translation inhibitors performed well in live mice, we set up a pulse-chase experiment with the harringtonine injection followed by the cycloheximide injection through the tail vein (Fig. 3A-B). The setup included two catheters connected to syringes filled with translation inhibitors on one side and inserted to each of the two major lateral tail veins on another side (Fig. 2A). The experiment was initiated by the injection of harringtonine and followed by the injection of cycloheximide after a specified time. We tested different intervals between harringtonine and cycloheximide injections, ranging from 15 to 120 sec (Fig. S1B), and found that 15, 30 and 45 sec time points were optimal. At first, we were surprised that the inhibitors affected translation so fast, much faster than in experiments involving cultured cells (4). While the *in vitro* experiment performed with a mouse cell line had an ~90 sec delay prior to harringtonine showing an effect, the ribosomal run-off was clearly observed already within 15 sec in live mice (Fig. 3E). On the other hand, we waited one minute after the injection of the second inhibitor and sacrificed the mouse by cervical dislocation. The heart was functioning normally pumping the blood. Thus, even if it has taken more time for both inhibitors to infuse organs and cells, the differential between injections has been kept constant.

**Fig. 3.**
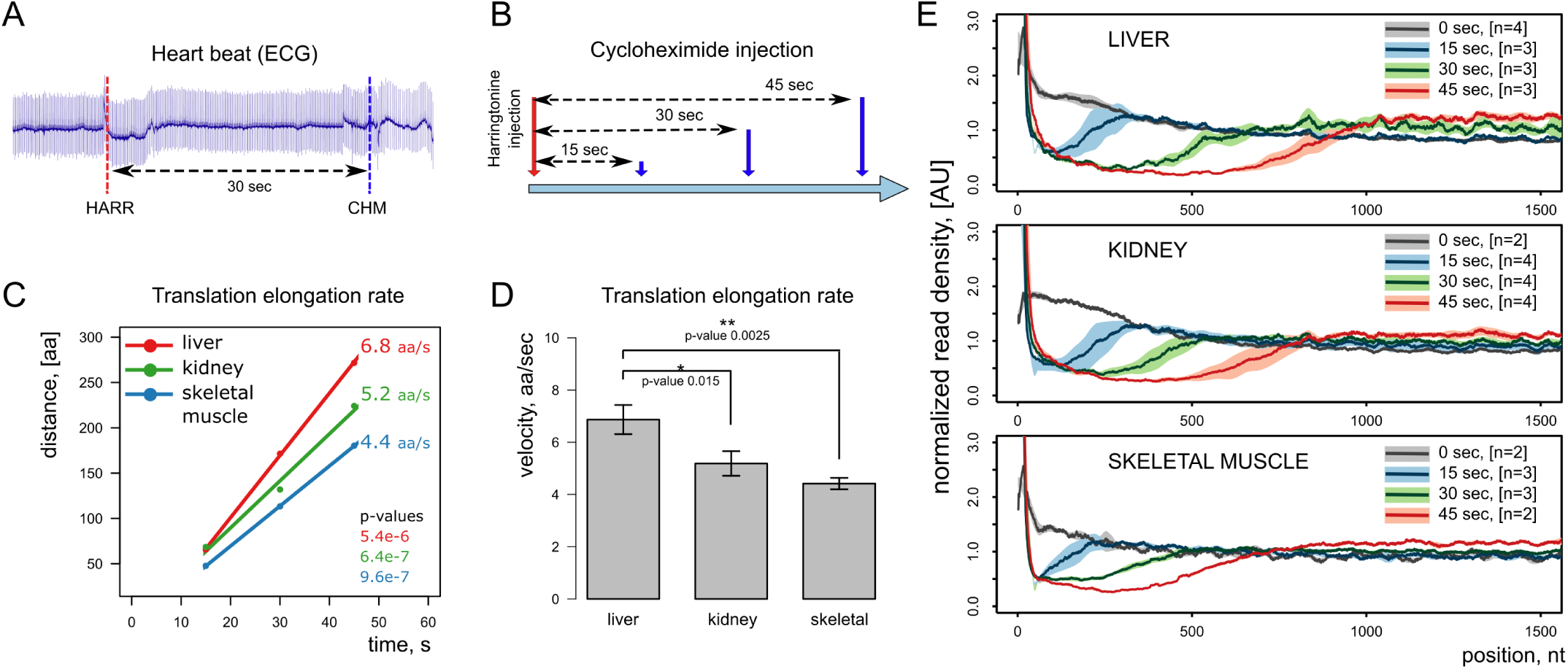
Measuring translation elongation rate *in vivo*. (A) ECG monitoring of heart function before and after drug injection to ensure the heart beat rate is unaffected at the chosen dosage. (B) Experimental design for a time-resolved study. (C) Linear regression analysis of injection timepoints to infer the elongation rate. (D) Comparison of mean translation elongation rates in three organs. (E) Metagene analysis of ribosome profiling sequences. Genes longer than 2,000 nucleotides are presented. The first 1,500 nucleotides are shown starting from the first codon of the reading frame. Solid lines represent mean signal from biological replicates, and light shading corresponds to a standard deviation.

To precisely measure translation elongation rate, we used 3-4 animals per time point, three time points in total. The resulting ribosome profiling libraries from three organs (live, kidney, skeletal muscle) were sequenced. Fig. 3E shows the cumulative ribosome coverage of open reading frames in these samples. Time dependent run-off was observed in every organ, and, interestingly, the rate was different between the organs. Linear regression analysis of ribosome coverage tracks revealed the translation rates of 6.8, 5.2, 4.4 amino acids per sec for liver, kidney and skeletal muscle, respectively (Fig. 3C-E). The rate difference may reflect the metabolic load of these organs. These rates can be compared with that previously estimated in mammalian cell culture (∼5.5 aa/sec) by Ribo-seq (4, 14), and in rat liver (~5.7 aa/sec) by radioactive labeling (9, 15).

### Selection of proper time points and the injection route is critical to accurately measure the translation elongation rate

As discussed above, the original 5 min long incubations with harringtonine resulted in sharp peaks at the translation start sites. However, ribosomes were still very abundant at the distant downstream regions of mRNAs in the liver as can be seen at the metagene level (Fig. 1C) as well as in individual gene cases (Fig. 4). This effect appeared exclusively in the liver and, at first, we hypothesized that it likely happened due to the lower blood pressure in the retro-orbital sinus and thus less efficient spread of translation inhibitors by the vascular system. However, the change of the injection route to the tail vein proved it wrong, i.e. a similar outcome was observed when injections were performed through the tail vein. Furthermore, given more timepoints and better time resolution, we encountered a sudden drop in the observable translation elongation rate past 45 sec not only in the liver but in other organs too. When 60 sec time points were sequenced and analyzed for liver, kidney, and skeletal muscle, all samples showed a decline in the observed translation rate compared to the 15-45 sec interval (Fig. S4). No retardation of the translation elongation rate was previously observed in a study involving mouse cell culture, where the first timepoint was at 90 sec (4). Similarly, no translation shutdown was reported when radioactive amino acids were used to estimate the elongation rate in rat and toadfish livers (8, 9, 15). This observation points to an organ-specific mechanism monitoring translation and shutting it down at the elongation stage when something goes wrong.

**Fig. 4.**
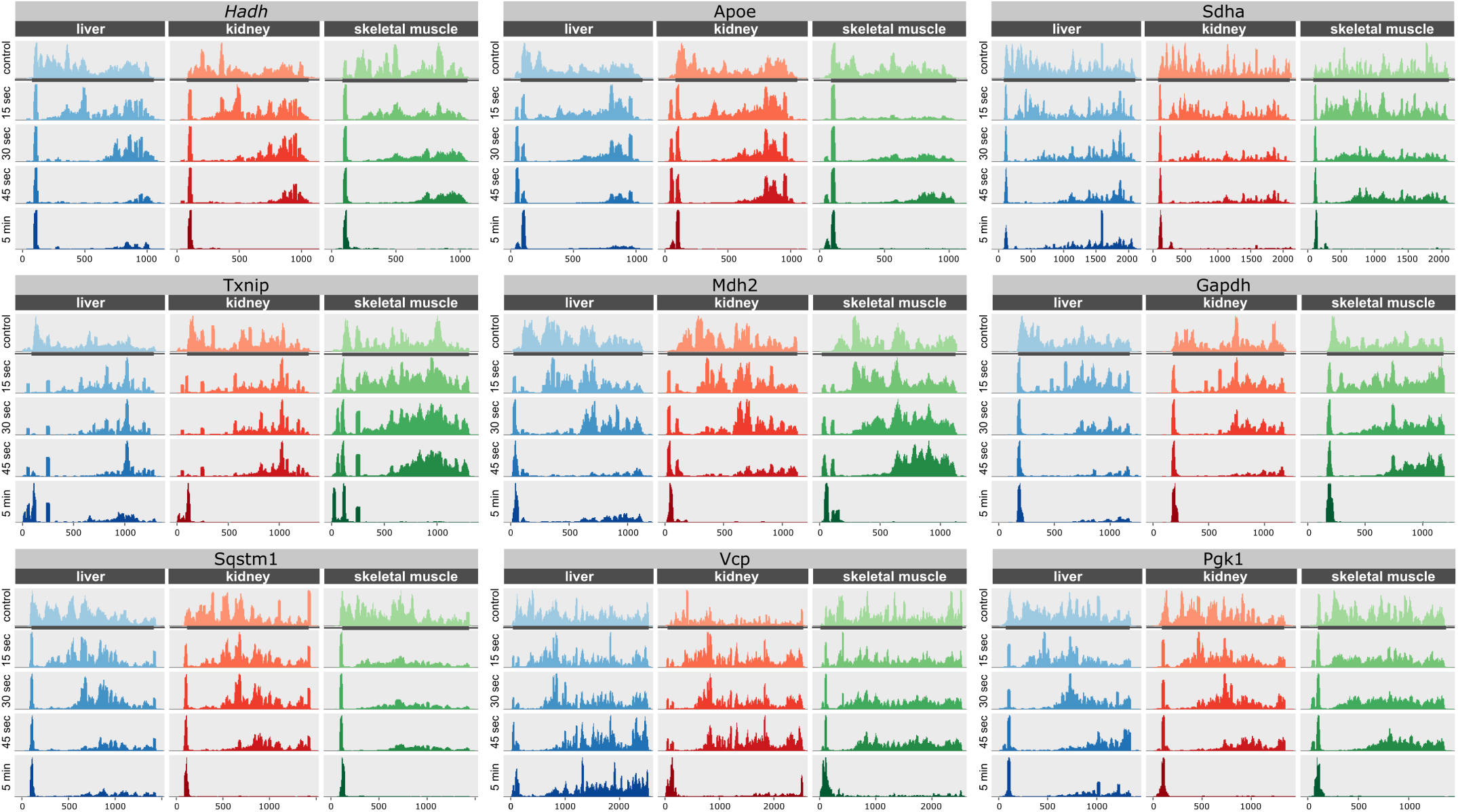
Ribosome footprint profile of representative transcripts. Transcripts were selected among those highly expressed in three organs: liver, kidney and skeletal muscle. For each transcript, the entire ORF was selected with extra 100 nucleotides from 5’- and 3’-UTRs. The ribosomal run-off is a consequence of the harringtonine injection. The entire ORF with 100 nucleotides from the 5’ and 3’ UTRs is shown. The sharp peak on the left side of each plot corresponds to ribosomes stalled at the translation start site by harringtonine. Annotated ORF is marked with the gray rectangle.

The fast response rate indicates the involvement of post-translational regulation, presumably a modification of elon-gation factors. Alternatively, the cellular concentration of harringtonine may reach the level inhibitory to elongation, since we administered way more drug compared to the *in vitro* study (7). It could also explain why the liver is the most affected organ in our set. Unlike heart and skeletal muscle, liver has sinusoidal (discontinuous) capillaries with large gaps in the basal membrane and between epithelial cells (16). It facilitates the exchange of larger biomolecules between blood and hepatocytes and increases the diffusion rate in general. Therefore, if harringtonine can interfere with elongation at high cytoplasmic concentrations, it should affect the liver in the first place. Investigation of the mechanism involved is a goal of our future studies.

In addition to the translation elongation rate, we could assess translation initiation *in vivo*, without prior cell extraction and thus without perturbing physiological conditions. In this regard, it provides a viable alternative to current approaches (17, 18). Ribosomes accumulate at the translation start sites over time after the harringtonine injection. This applies both to the classical open reading frames (Fig. 4) and regulatory upstream reading frames as can be seen in the case of the *ATF4* transcript - the well-known example of translational regulation mediated by upstream ORFs (Fig. 5).

**Fig. 5.**
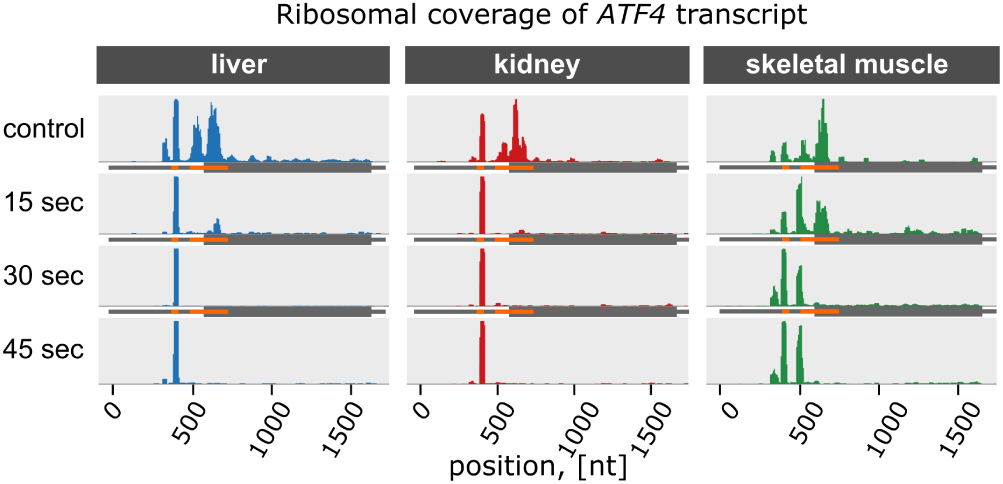
Ribosome footprint coverage of ATF4 transcript. This gene is a classic example of post-transcriptional regulation mediated by upstream reading frames (uORFs). It has 2 uORFs, shown in orange. Shortly after the injection of harringtonine, ribosomes are stalled at the start codons of the corresponding uORFs. Interestingly, there are two peaks in skeletal muscle samples: one at the first uORF and another at the second. However, there is only a single peak in liver and kidney.

## Measuring translation efficiency across multiple organs

We further complemented ribosome profiles with transcriptomes from the same organs. The ratio of Ribo-seq to mRNA-seq counts measures translation efficiency. It reflects how well the transcript is translated compared to other transcripts from the same organ. Transcripts with higher translation efficiency yield more protein molecules given the same number of transcripts. Calculating this ratio for every single transcript gives us the list of organ-specific translation efficiencies for thousands of proteins. Gene set enrichment analysis revealed that pathways involved in metabolism were enriched in genes with higher translation efficiency. These pathways included TCA cycle, oxidative phosphorylation, fatty acid metabolism, and glycolysis. On the other hand, mTOR signaling, MAP kinase signaling, insulin signaling and ribosome (as a GO term) were enriched with genes with lower translation efficiency (Fig. S5).

## Discussion

Development of the method for ribosome profiling *in vivo* allowed us to determine translation elongation rate in multiple organs of mice. These rates significantly different among organs, with liver being fastest in synthesizing proteins, and skeletal muscle being slowest among. One concern we had prior to this study is the binding properties of albumin. It contains multiple hydrophobic binding pockets to retain and transport various fatty acids, steroids, and drugs. In a similar time-course experiment in cell culture, harringtonine was mostly bound by albumins coming from the bovine serum supplement. For this reason, we injected very high amounts of harringtonine and cycloheximide in the bloodstream. Although it is currently difficult to estimate the dynamics of translation inhibitor uptake by cells *in vivo*, our results indicate that the minimal effective drug concentration is likely to be much lower than what was used in the study. It is also possible that a lower amount of harringtonine might not interfere with elongation at late time points (above 45 sec).

Our approach allows targeting multiple organs, and a mouse providing a single time point for up to 8 organs. However, experiments can be easily modified to produce multiple time points for a single organ. Translation inhibitors are well suited for perfusion-like procedures, when the major blood vessels of an organ, for instance, the liver, is surgically accessed and the organ is perfused or washed with a drug-containing buffer. Unlike radioactive labeling, small molecule drugs used in animal studies possess no harm to experimentalists, and the collected tissues could be used for transcriptome sequencing, and proteomic and metabolomic studies. It also expands the existing puromycin-based technique for spatiotemporal monitoring of nascent peptide release (19).

We explored the feasibility of measuring translation elon-gation in multiple organs and found that it is possible even within very narrow time intervals. Due to a broad selection of organs to test, we limited the number of time points to 3. While it was sufficient to estimate the mean translation rate with high precision, gene-specific translation rates require better temporal resolution. The method presents new opportunities for animal studies and may be amenable to study the effects of ribosome protein mutations, dietary interventions, age or other factors on translation.

Concluding, our method allows measuring the elongation rates in multiple organs and cell types *in vivo*. In addition, it captures the profiles of translation start sites, upstream reading frames and organ-specific translation efficiency. Elon-gation rates may be influenced by multiple factors, such as post-translational modifications of ribosomal proteins, local levels of amino acids, and tRNA concentrations. The finding of the efficient delivery of translation inhibitors to mouse organs would allow the use of additional effectors, targeting termination and various mechanisms of regulation of protein synthesis. Proteostasis is known to be associated with physiological conditions such as aging (20), cancer (21), and ribosomal pathologies (22). Our study lays out a path for interrogating translation directly in animal models in the unperturbed setting.

## Materials and Methods

### Translation inhibitors used in experiments

Lactimidomycin was purchased from EMD Millipore. Harringtonine was available from three vendors: Abcam, Santa Cruz Biotechnology, and Carbosynth. We tested all three sources and found no difference in quality and toxicity. Most of the experimental work was done with the chemical supplied by Abcam. Cycloheximide was purchased from Sigma-Aldrich.

### Drug delivery in mice

We used two injection routes: a tail vein and a retro-orbital sinus. Retro-orbital injection requires little practice and is convenient for single drug delivery (12). It is quite invasive, therefore, we sedated mice with the intraperitoneal injection of pentobarbital (20 mg/kg). Tail veins are more appropriate for accurately timed double injections. There are two large lateral veins in mouse tail. We attached catheters to both veins and connected them to syringes filled with harringtonine or cycloheximide. During the entire procedure, mice were sedated with a continuous flow of 1% isoflurane mixed with oxygen (Visualsonics, Vevo anesthesia system). Sedation caused no significant change in translation based on polysome analyses. In a typical experiment, we injected 200 *μ*l of harringtonine (5 mg/ml) in phosphate buffer saline (PBS), followed by 100 *μ*l of cycloheximide (20 mg/ml) in the same buffer. Cycloheximide is highly soluble in the buffer. Harringtonine, on the other hand, is poorly soluble in aqueous solutions; therefore, 50 mg/ml stock was prepared in dimethyl sulfoxide and 20-fold dilution in phosphate buffer saline was made fresh right before the injection. Lactimidomycin was prepared in the similar fashion. The highest dose of lactimidomycin that we could administer was 200 *μ*l of a 0.25 mg/ml in 20% dimethyl sulfoxide, 80% PBS. Several attempts were made to increase the amount of harringtonine cycloheximide that could be deliver to a mouse. Supplementing PBS with 40% (2-hydroxypropyl)-*β*-cyclodextrin (Sigma) increased cycloheximide solubility up to 60 mg/ml without any toxic response from mice. It did not help with harringtonine or lactimidomycin solubility. Heart function was monitored by ECG (AD Instruments, Powerlab 8/30 recorder with BioAmp electrodes). Animals with severe drop of heartbeat or incomplete injection were excluded from the analysis. All mice used in this study were C57BL/6j males, 10-15 weeks of age, from Jackson Lab.

### Harvesting tissues

Unless otherwise stated, in the case of single inhibitor injection, we held a mouse asleep for 5 min before euthanizing it by the cervical dislocation. Mice with time-course injections of two inhibitors were kept alive for 1 min after the second inhibitor injection, then euthanized by the cervical dislocation. Nine organs were collected (liver, kidney, lungs, heart, skeletal muscle (lower limbs), spleen, pancreas, testes, and brain). Large organs were cut in slices, snap frozen in liquid nitrogen and stored at -80°C.

### Ribosome profiling

In a typical experiment, we used 30 mg of the liver, 60 mg of kidney and 200 mg of skeletal muscle to extract ribosomes. Soft tissues, such as liver, kidney, pancreas, and brain were lysed in a glass-Teflon Dounce homogenizer filled with the ice-cold buffer. Tough tissues, such as heart, skeletal muscles and lungs were first pulverized in a ceramic mortar-pestle filled with liquid nitrogen, then lysed a glass-Teflon Dounce homogenizer. Based on our previous experience, RNase S7 and T1 are the best ribonucleases for ribosome footprint generation in mouse organ lysates (5). Lysates from control, non-injected and harringtonine-injected mice were treated with the 4:1 mixture of RNase T1 (Epicentre, cat# NT09500K) and RNase S7 (Roche/Sigma, cat# 10107921001). RNase S7 comes as powder and the 10 mg/ml stock was made in 10 mM Tris-HCl pH 7.5, 50 mM NaCl, 1 mM EDTA, 50% glycerol. Tissue lysates from mice used in time-course experiments were treated with RNase T1 alone. Different organs required specific lysis conditions. Lysis buffer composition for liver and kidney was as follows: 20 mM Tris-HCl pH 7.5, 100 mM KCl, 5 mM MgCl2, 1 mM DTT, 1% Triton, 0.1 mg/ml cycloheximide. Lysis buffer composition for skeletal muscle: 20 mM Tris-HCl pH 7.5, 100 mM KCl, 5 mM MgCl2, 1 mM DTT, 1% Tween-20, 0.25% deoxycholate, 0.1 mg/ml cycloheximide. Lysis buffer composition for lung, pancreas, spleen, heart, brain, and testis: 20 mM Tris-HCl pH 7.5, 100 mM KCl, 5 mM MgCl2, 1 mM DTT, 1% Tween-20, 0.25% deoxycholate, 0.1mg/ml cycloheximide. For preparative purposes, i.e. when ribosomes were used for footprint extraction and sequencing, we kept 5 mM MgCl2 in lysis buffers. For quality checks and sucrose gradient analyses, we increased MgCl2 up to 10 mM. Lysis buffers were also supplemented with protease inhibitors (Roche) and 5 mM CaCl2 as RNase S7 was applied. Sucrose gradients were always 10-50% sucrose in 20 mM Tris-HCl pH 7.5, 100 mM KCl, 10 mM MgCl2, 1 mM DTT, 0.1 mg/ml cycloheximide. Fractionation was performed by ultracentrifugation for 3 h at 35,000 rpm in an SW41 rotor (Beckman, Optima L-20K) at 4°C. After the centrifugation, gradients were passed through a UV detector (Bio-Rad) and the absorption at 254 nm was recorded. The fraction containing monosomes was collected in a single tube if necessary. The volume of the sample was brought to 50 *μ*l by concentrating it using 100 kDa filters (Amicon Ultra, Millipore). Then, the sample was diluted to 500 *μ*l with a buffer containing 10 mM Tris-HCl pH 7.5, 2 mM EDTA, 1% SDS. RNA was extracted by hot acid phenol (Ambion) and precipitated by the glycogen-ethanol method (1/10 volume of 3 M sodium acetate, 1/100 volume of glycogen, 2.5 volumes of pure ethanol, 1-hour incubation at -20°C followed by centrifugation). RNA was loaded on a 15% polyacrylamide TBE-urea gel and the band containing ribosomal footprints around 28 nucleotides was cut. Subsequent steps were the same as described in (5).

### Transcriptome library preparation

Total RNA was extracted from tissues with Trizol (Ambion) and purified using Direct-zol 96-well plate RNA kit (Zymo Research). Sample quality was checked with TapeStation (Agilent). Every tissue except pancreas had the RNA integrity number (RIN) higher than 8.7 which indicates high-quality RNA. Extraction of intact RNA from the pancreas is hardly possible due to the presence of the endogenous RNase A in high amounts, therefore we proceeded with sequencing although the RIN was *∼*6.

### High throughput sequencing and data processing

Ribosome profiling libraries were sequenced on the Illumina HiSeq 2000 and NextSeq 500 platforms at the Harvard University Bauer Core. mRNA-seq libraries were sequenced on the Illumina HiSeq 4000 in the paired end mode at the Novogene Inc. Adapters were removed from ribosome profiling reads with Cutadapt software (cutadapt -u 1 -m 23-a AGATCGGAAGAGCACACGTCT) (23). mRNA-seq sequences were aligned with TopHat 2.1.0 using following settings: tophat –transcriptome-index –no-discordant –no-mixed –no-novel-juncs (24).

The NCBI mouse genome build GRCm38.p3 and the Mus musculus Annotation Release 105 were used as a reference. When aligning mRNA-seq and ribosome profiling reads for expression and translation efficiency estimation we used the following strategy. Only full chromosomes were left and all non-chromosomal and mitochondrial records removed. Furthermore, only RefSeq and BestRefSeq records were left in the transcriptome annotation, while Gnomon predictions were discarded along with pseudogenes. Read count per gene was accessed by HTseq-count software (25).

To plot time-dependent ribosome occupancy (Fig. 3E), we employed a different strategy. First, using a gene bank (gbk) file, which comes in a package with the genome assembly and annotation, we extracted RefSeq and BestRefSeq records for every gene including CDS, 5’-UTR and 3’-UTR lengths and sequences. Among them, we identified the longest isoform for every gene, prioritized as CDS > 5’UTR > 3’UTR. 5’-UTRs were trimmed by 100 nucleotides. If either or UTRs were shorter than 100 nucleotides, we filled it with up to 100 based on genomic coordinates. To prepare a list of non-redundant genes, we run blast of all vs. all (blastall -p blastn-m 8 -b 500 -v 500 -e 0.001). Gene pairs that are too similar at the level of nucleotide sequence were excluded. In addition to the e-score, we enforced a requirement of the high-homology stretch being at least 50 nt long, and if it was longer, the similarity had to be at least 90% to treat these genes as homologous and redundant. A total of 13,685 genes passed every threshold. This reference set ensured unambiguous alignment of ribosome footprints.

### Gene set enrichment analysis

To identify pathways associated with genes with significant high or low translation efficiency across tissues, we used *limma* software (26) to calculate p-value for each gene against the null hypothesis that average translation efficiency of the gene is equal to the average translation efficiency across whole transcriptome. When several replicates of the same tissue were present, we considered translation efficiency of every gene for the corresponding tissue as an average across all replicates. Then we performed a pathway GSEA on a pre-ranked list of genes (27). For every gene, this list contained z-score, calculated as: *ln*(*P*) *S* where P is the p-value calculated as described previously and S is the sign of difference between average TE of the gene and average TE across whole transcriptome. Kyoto Encyclopedia of Genes and Genomes (KEGG), Reactome, Biocarta and Gene Ontology (GO) Biological Process (BP) and Molecular Function (MF) databases were used in this analysis.

### Cluster analysis

Translation efficiency (TE) for individual genes were calculated as Ribo-seq read counts divided by mRNA-seq counts. Only genes expressed in all organs were taken into analysis (>= 10 reads in every organ and sequencing type, 4782 genes satisfied these criteria). Normalization across organs were performed by centering TE values distribution at 0 and dividing by the standard deviation. Organs were hierarchically clustered by complete linkage method using Pearson correlation distance (1 - *cor*). Genes were hierarchically clustered by Ward method using Euclidean distance. The genes dendrogram was further sorted by dendsort R package with default settings (28).

Experimental animal protocols were approved by the Institutional Animal Care and Use Committee at the Brigham and Women’s Hospital, Harvard Medical School.

## ACKNOWLEDGMENTS

We thank Margarita Meer, Veronica Silva and staff of the Animal Surgery and Physiological Core at Brigham and Women’s Hospital for assistance with tail vein injections and heart function monitoring. We also thank Alexander Tyshkovskiy for guidance in GSEA analysis. Supported by NIH DK117149.

